# FSTest: an efficient tool for cross-population fixation index estimation on variant call format files

**DOI:** 10.1101/2022.08.16.503949

**Authors:** Seyed Milad Vahedi, Siavash Salek Ardestani

**Author notes:** **Corresponding authors:** Seyed Milad Vahedi, MSc, DVSc, Department of Animal Science and Aquaculture, Dalhousie University, Bible Hill, NS, B2N5E3, Canada.

## Abstract

Fixation index (Fst) statistics provide critical insights into the evolutionary processes affecting the structure of genetic variation within and among populations. The Fst statistics have been widely applied in population and evolutionary genetics to identify genomic regions targeted by selection pressures. The FSTest 1.3 software was developed to estimate four Fst statistics of Hudson, Weir and Cockerham, Nei, and Wright using high-throughput genotyping or sequencing data. Here, we introduced FSTest 1.3 and compared its performance with two widely used software of VCFtools 0.1.16 and PLINK 2.0. Chromosome 1 of 1000 Genomes Phase III variant data belonging to South Asian (N = 211) and African (N = 274) populations were included as an example case in this study. Different Fst estimates were calculated for each single nucleotide polymorphism (SNP) in a pairwise comparison of South Asian against African populations, and the results of FSTest 1.3 were confirmed by VCFtools 0.1.16 and PLINK 2.0. Two different sliding window approaches, one based on a fixed number of SNPs and another based on a fixed number of base pair (bp) were conducted using FSTest 1.3 and VCFtools 0.1.16. Our results showed that regions with low coverage genotypic data could lead to an overestimation of Fst in sliding window analysis using a fixed number of bp. FSTest 1.3 could mitigate this challenge by estimating the average of consecutive SNPs along the chromosome. FSTest 1.3 allows direct analysis of VCF files with a small amount of code and can calculate Fst estimates on a desktop computer for more than a million SNPs in a few minutes. FSTest 1.3 is freely available at https://github.com/similab/FSTest.

## Introduction

Fixation index (Fst) includes a group of statistics that quantify levels of genetic differentiation between populations based on allele frequencies (Biswas and Akey, 2006). Fst can be used to identify loci at which a beneficial allele has recently gone to fixation (Holsinger and Weir, 2009). The Fst has been conceptually defined in many ways (Hudson *et al*., 1992; Wright, 1949), among which the Weir and Cockerham (1984) and Nei (1973) estimations are the most popular statistics. Two widely used software, including VCFtools 0.1.16 (Danecek *et al*., 2011) and PLINK 2.0 (Purcell *et al*., 2007), have provided the functions to compute the Fst values; the first one estimates Weir and Cockerham’s and the latter estimates Weir and Cockerham’s and Hudson’s.

Fst values can be estimated either per single nucleotide polymorphism (SNP) or as an average per genomic windows harboring multiple SNPs, referred to as “sliding window” analysis. In the sliding window analysis, Fst statistics are calculated for a small window of the data; however, windows can be determined based on a fixed number of base pairs (bp) or a fixed number of SNPs. While VCFtools provides sliding window modes only based on bp, PLINK lacks this feature. However, sliding window analysis on a fixed number of bp, particularly when low coverage genotypic data is utilized, leads to overestimation or underestimation of statistics since the number of SNPs inside windows would be inconsistent (Gusnanto *et al*., 2014). Therefore, a tool that can implement sliding window analysis using a fixed number of SNPs could be an asset. Moreover, except for VCFtools, a limited number of tools are currently available to produce Fst statistics from variant call format (VCF) data. Here, we introduced FSTest 1.3 software, a single package that completely covers well-known estimates of Fst, performs both SNP-based and sliding windows analyses, and supports VCF files as inputs. The results of the software were compared with those obtained from VCFtools 0.1.16 and PLINK 2.0.

## Materials and methods

FSTest 1.3 is implemented in Python and accepts VCF files as input. The tool automatically manages missing genotypes and supports phased and unphased genotypic data. FSTest calculates between-population Fst estimators of Hudson *et al*. (1992), Weir and Cockerham (1984), Nei (1973), and Wright (1949) as follows:

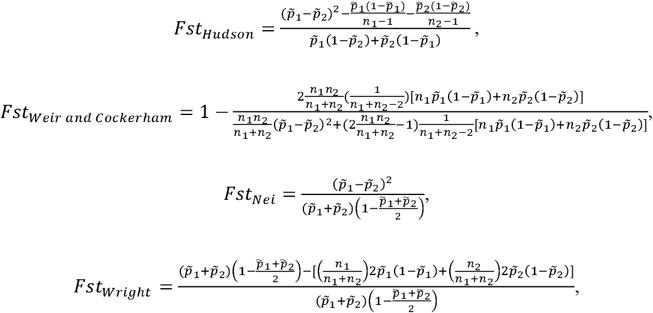

where *m* is the sample size, and 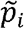 is the sample allele frequency in population *i* for *i* ∈ {1, 2}. All Fst estimations range between 0 and 1, and values less than zero are denoted zero by the software as they do not have a biological interpretation.

For sliding window analysis, the average Fst values across SNPs will be taken for each window. Users can also implement Z-transformation on all Fst estimates for SNP- or window-based approaches to get unbiased Z_Fst_ values as given:

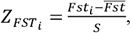

where *Fst*_*i*_ is the Fst value of *i*th SNP or *i*th window, 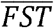 and *S* represent the average value and standard deviation of Fst between two populations estimated from all SNPs or windows. We have also added the functionality of creating publication-quality Manhattan plots for the results of SNP and sliding window analyses.

Here, we demonstrated an example case by analyzing chromosome 1 of 1000 Genomes Phase III variant data downloaded from the project’s FTP site (https://ftp.1000genomes.ebi.ac.uk/). A total of 485 samples belonging to two different ancestries, South Asian (N = 211) and African (N = 274), were included in this study. Quality control was performed on 6,468,094 SNPs on chromosome 1 using PLINK 2.0 software (Purcell *et al*., 2007). Markers with minor allele frequency□< □0.01, SNP calling rate □< □0.90, and extreme departure from Hardy–Weinberg equilibrium (p-value□< □10^−7^) were removed. Individuals with missing genotype data > 0.1 were also removed from the study. After quality control, a total of 1,206,810 SNPs from 485 individuals remained for further analysis. The average±standared error (SE), minimum, and maximum distance between SNPs in the final genotypic data were 207±17, 1, and 21,050,296 bp. The longest distance between SNPs belonged to the region of 12,148,5143-142,535,439 bp.

Fst values of Hudson, Weir and Cockerham, Nei, and Wright were estimated using FSTest 1.3, Hudson’s and Weir and Cockerham’s using PLINK 2.0 (Purcell *et al*., 2007), and Weir and Cockerham’s by VCFtools 0.1.16 (Danecek *et al*., 2011) (Table 1). Spearman’s correlations among the estimates were calculated, and a correlogram was constructed using corrplot 0.92 package (Wei *et al*., 2017) in R 4.1.0 (R Core Team, 2021).

**Table 1.** Different features of Fixation index (Fst) estimation using FSTest 1.3, VCFtools 0.1.16, and PLINK 2.0 software. Different Fst values per SNP along with two different window-based approaches, in which the window is determined based on a fixed number of base pairs (Win-bp) or a fixed number of SNP (Win-SNP), were compared among the tools.

| Software | Fst estimates | Approach |  |  |
| --- | --- | --- | --- | --- |
|  |  | SNP | Win-SNP | Win-bp |
| **FSTest 1.3** | $F_{st_{Hudson}}$, $F_{st_{Weir}}$ and $Cockerham$,<br>$F_{st_{Wright}}$, $F_{st_{Nei}}$ | ✓ | ✓ | - |
| **VCFtools 0.1.16** | $F_{st_{Weir}}$ and $Cockerham$ | ✓ | - | ✓ |
| **PLINK 2.0** | $F_{st_{Hudson}}$, $F_{st_{Weir}}$ and $Cockerham$ | ✓ | - | - |

Two sliding window approaches, one based on a fixed number of SNPs and another based on a fixed number of bp, were applied using FSTest 1.3 and VCFtools 0.1.16 software, respectively (Table 1). In the first one, a 500 SNPs sliding window with a step size of 50 SNPs, and in the latter, a 100 kb sliding window with a step size of 10 kb was used. PLINK 2.0 software was not included in the window-based analyses as it lacks this feature.

A desktop computer with AMD Ryzen 7 4800H (8-core) CPU and 32 gigabytes installed RAM was used for all analyses. The time of analyses was measured using the log files and compared among the software. CMplot package (Yin *et al*., 2021) in R 4.1.0 (R Core Team, 2021) was used to create circos and Manhattan plots. Visualization of chromosomal segments harboring *Fst*_*mean*_ > 0.15 was performed using ChromoMap 0.4.1 (Anand and Rodriguez Lopez, 2022) in R 4.1.0 (R Core Team, 2021).

## Results and discussion

Different Fst statistics were estimated for 1,206,810 SNPs in the pairwise comparison of chromosome 1 between South Asian and African populations (Fig. 1a). Estimates of *Fst*_*Hudson*_ and *Fst*_*weir and cockerham*_ were comparable among the results of FSTest 1.3, VCFtools 0.1.16, and PLINK 2.0 (Table 2). We obtained similar means±SE for *Fst*_*Hudson*_ and *Fst*_*weir and cockerham*_ (0.07±0.00), as well as for *Fst*_*wright*_ and *Fst*_*Nei*_ (0.04±0.00). The highest and the lowest range of Fst values belonged to the Nei (0-0.85) and Wright (−0.08-0.39) estimators, respectively. Spearman’s correlations between different Fst estimates were obtained, and a correlogram was constructed (Fig 1b). We observed 100% correlations between *Fst*_*Hudson*_ and *Fst*_*weir and cockerham*_ estimates achieved by FSTest 1.3 and those by VCFtools 0.1.16 and PLINK 2.0 (Table 2). Descriptive statistics and correlations of FSTest 1.3 results with those from VCFtools 0.1.16 and PLINK 2.0 confirms the accuracy of FSTest 1.3 performance.

**Table 2.** Mean, median, standard error (SE), and range of different Fst estimates per SNP.

| **Tool** | **Statistics** | **Mean** | **Median** | **SE** | **Range** |
| --- | --- | --- | --- | --- | --- |
| **FSTest1.3** | $Fst_{Hudson}$ | 0.07 | 0.04 | 0 | 0-0.59 |
| | $Fst_{Weir\ and\ Cockerham}$ | 0.07 | 0.04 | 0 | 0-0.56 |
| | $Fst_{Wright}$ | 0.04 | 0.02 | 0 | -0.08-0.39 |
| | $Fst_{Nei}$ | 0.04 | 0.04 | 0 | 0-0.85 |
| **VCFtools 0.1.16** | $Fst_{Weir\ and\ Cockerham}$ | 0.07 | 0.04 | 0 | 0-0.56 |
| **PLINK 2.0** | $Fst_{Hudson}$ | 0.07 | 0.04 | 0 | 0-0.59 |
| | $Fst_{Weir\ and\ Cockerham}$ | 0.07 | 0.04 | 0 | 0-0.56 |

**Figure 1.**
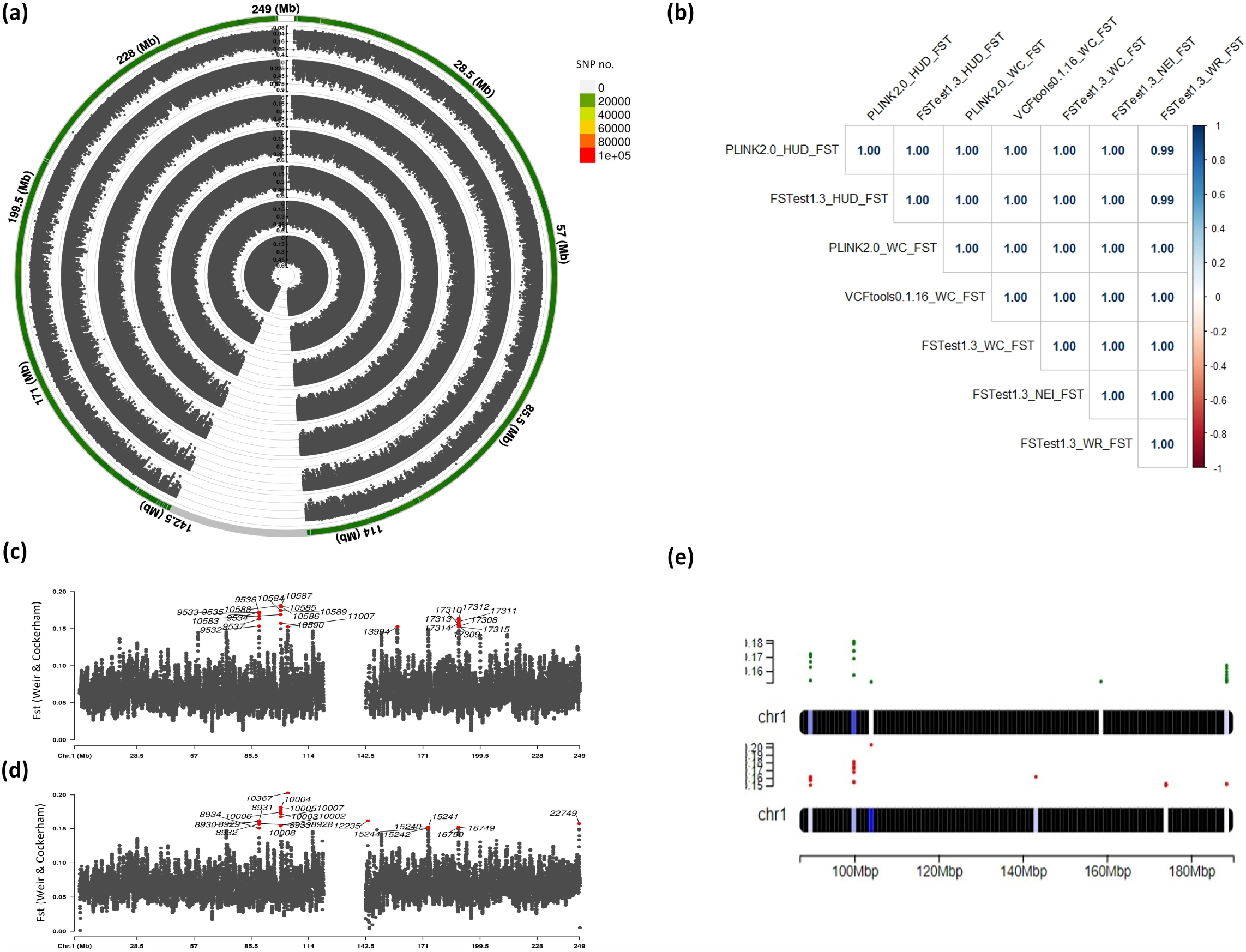
Circos plot (a) shows different Fst values estimated by FSTest 1.3, PLINK 2.0, and VCFtools 0.1.16 tools. From inside to outside, layers represent the distribution of *Fst*_*Hudson*_ obtained from PLINK 2.0, *Fst*_*Hudson*_ from FSTest 1.3, *Fst*_*weir and cockerham*_ from PLINK 2.0, *Fst*_*weir and cockerham*_ from VCFtools 0.1.16, *Fst*_*weir and cockerham*_ from FSTest 1.3, *Fst*_*Nei*_ from FSTest 1.3, and *Fst*_*wright*_ from FSTest 1.3. Correlogram (b) demonstrates the Spearman’s correlations among different Fst estimations by applied tools. Manhattan plots (c) and (d) show the distribution of *Fst*_*weir and cockerham*_ using two sliding window analyses by FSTest 1.3 and VCFtools 0.1.16. A sliding window of 500 SNPs with a step size of 50 SNP was applied to FSTest 1.3, and a sliding window of 100kb with a step size of 10 kb was implemented in VCFtools 0.1.16. Red points represent windows with *Fst*_*mean*_ > 0.15 and numbers show the number of windows. Chromosome visualizations (e) and (f) depicts the overlapping windows with *Fst*_*mean*_ > 0.15 obtained by FSTest 1.3 and VCFtools 0.1.16, respectively.

*Fst*_*weir and cockerham*_ was estimated using two different sliding window approaches: (i) based on a fixed number of SNP using FSTest 1.3, and (ii) based on fixed number of bp using VCFtools 0.1.16, and the results were shown in Fig 1c and Fig 1d, respectively. To compare the results of two approaches, windows with *Fst*_*mean*_ > 0.15 were highlighted (Fig 1c and 1d) and overlapping segments were compared (Fig 1e). We identified 24 and 23 genomic windows with *Fst*_*mean*_ > 0.15 using FSTest 1.3 and VCFtools 0.1.16, respectively. As shown in Figure 1e, among FSTest 1.3 results, the highest Fst values belonged to the region of 100,000-102,500 kb, containing eight genomic windows with *Fst*_*mean*_ > 0.15 (*Fst*_*mean*_ = 0.17). In contrast, this region was ranked second by VCFtools 0.1.16, after the 102,500-105,000 kb segment, which only contained one genomic window with *Fst*_*mean*_ > 0.15. After checking the results of VCFtools 0.1.16, we realized that this genomic window only harbors one SNP. We also found that sliding window analysis using a fixed number of bp might be more sensitive to the edges of genomic segments. Fig 1d demonstrates that one of the windows with *Fst*_*mean*_ > 0.15 (no. 12235; chr1: 143,540-143,640 kb) was located after the segment without variants in the middle of chromosome 1 and only contained 10 SNPs. Our results are in parallel with Gusnanto *et al*. (2014) findings that regions with low coverage genotypic data could lead to overestimation or underestimation of Fst as the number of SNPs inside windows would be inconsistent. Therefore, the interpretation of the VCFtools 0.1.16 results should always be with attention to the number of SNPs in the genomic windows. Meanwhile, FSTest could tackle this problem by estimating the average of consecutive SNPs along the chromosome.

Regarding the analysis runtime, FTSest 1.3 performed slightly slower than VCFtools 0.1.16 (Table 3). Reading VCF files can be time-consuming, mainly when dealing with genomic data from whole-genome sequencing or large-scale studies. VCF files contain structured data with numerous fields and annotations for each variant, such as variant type, quality scores, and genotype information, which must be processed. To mitigate the time-consuming nature of reading VCF files, parallel processing techniques can help improve the speed of reading and processing VCF files, which is suggested for the next versions. PLINK 2.0 performed significantly faster than FSTest 1.3 and VCFtools 0.1.16 due to using light PLINK format files as

**Table 3.**
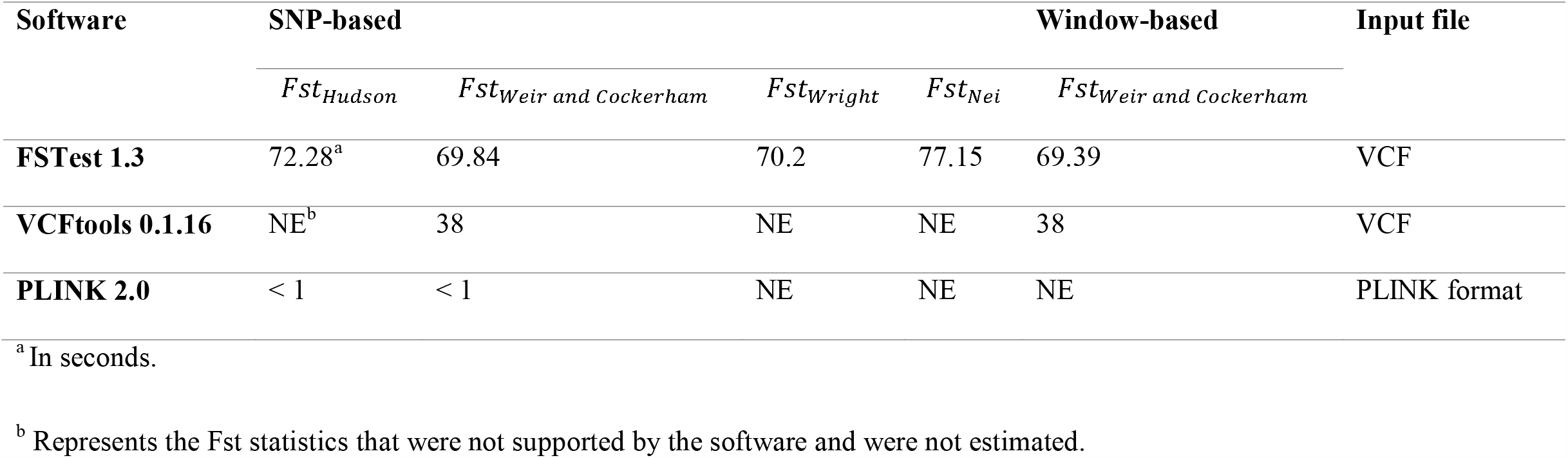
Comparison of runtime among different tools and Fst estimates included in this study.

In conclusion, FSTest 1.3 software contains extra pairwise Fst estimation methods, which are not present in the VCFtools 0.1.16 and PLINK 2.0 software. The tool allows direct analysis of VCF files with a small amount of code and is able to conduct sliding window analyses based on a fixed number of SNPs. FSTest 1.3 can calculate Fst estimates the results on a desktop computer for more than a million SNPs in a few minutes.

## Funding

We have received no specific funding for this study.

## Data availability

The genotypes used in this study is available at: https://ftp.1000genomes.ebi.ac.uk/

## Author contributions

SV and SS developed the original FSTest software. SV wrote the manuscript. All authors read and approved the final manuscript.

## Acknowledgments

We acknowledge the valuable comments provided by the reviewer, which greatly contributed to improving the quality of the paper.

## Notes

https://github.com/similab/FSTest

### Competing Interest Statement

The authors have declared no competing interest.

### Summary of Updates

FSTest 1.3 performance was compared with two widely used software of VCFtools 0.1.16 and PLINK 2.0.

